# Dynamic changes in somatosensory and cerebellar activity mediate temporal recalibration of self-touch

**DOI:** 10.1101/2023.12.01.569520

**Authors:** Konstantina Kilteni, H. Henrik Ehrsson

## Abstract

An organism’s ability to accurately anticipate the sensations caused by its own actions is crucial for a wide range of behavioral, perceptual, and cognitive functions. Notably, the sensorimotor expectations produced when touching one’s own body attenuate such sensations, making them feel weaker and less ticklish and rendering them easily distinguishable from potentially harmful touches of external origin. How the brain learns and keeps these action-related sensory expectations updated is unclear. We employed psychophysics and functional magnetic resonance imaging to pinpoint the behavioral and neural substrates of dynamic recalibration of expected temporal delays in self-touch. Psychophysical results revealed that self-touches were less attenuated after systematic exposure to delayed self-generated touches, while responses in the contralateral somatosensory cortex that normally distinguish between delayed and nondelayed self-generated touches became indistinguishable. During the exposure, the ipsilateral anterior cerebellum showed increased activity, supporting its proposed role in recalibrating sensorimotor predictions. Moreover, responses in the cingulate areas gradually increased, suggesting that as delay adaptation progresses, the nondelayed self-touches trigger activity related to cognitive conflict. Together, our results show that sensorimotor predictions in the simplest act of touching one’s own body are upheld by a sophisticated and flexible neural mechanism that maintains them accurate in time.

## Introduction

Our perception is shaped by the predictions we make about our self and the world and against which we compare our incoming sensations ^1–5^. One source of these predictions is based on the motor signals of our voluntary movements. Accordingly, the brain implements an internal forward model that represents our motor apparatus and the environment and that associates motor commands with their sensory consequences. Based on these associations, the brain forms predictions about the expected sensory consequences of a particular movement (*i.e.*, sensory feedback) including their timing, given a copy of the motor command (“efference copy”) ^6–10^. These predictions are essential; they enable compensate for neural delays in receiving and processing the actual sensory feedback ^11,12^, prospectively correct motor errors ^7^, improve the estimation of the current state of our body ^8,13,14^ and attenuate the received sensory feedback (*i.e.*, self-generated sensations) to increase the salience of externally generated sensations ^6,15–18^.

Nevertheless, although our perception is stable, the dynamics of our body and the world change. For example, as we grow, the neural delays in receiving and processing sensory feedback change ^19^. Similarly, throughout our life, we learn with practice to manipulate objects with novel dynamics that we do not know beforehand ^20^. Therefore, it is fundamental that our nervous system detects persistent errors between the expected and received sensory feedback of the movement (*i.e.*, prediction errors) and uses these errors to recalibrate the internal model to the new statistics, thereby ensuring accurate predictions, stable perception and adaptive control ^7,21–25^. To test for such recalibration, experimental studies have typically introduced perturbations between participants’ movements (*e.g.*, arm movements or speech) and the associated sensory consequences and tested whether participants’ behavior changes as the result of the adaptation to the perturbations ^7,23,26^. In the domain of time, several earlier studies have injected delays between the movement and the associated visual/auditory/vestibular consequences and showed that participants adapt to delays by adjusting their movements and/or perceptual biases ^27–38^. Importantly, these effects become much smaller or negligible in the absence of movement ^33,39^.

Regarding adaptation to delays in the somatosensory consequences of actions, Witney et al. (1999) showed that the timing of grip force responses typically observed when an object we hold in a precision grip is pulled on by our other hand ^41,42^ shifted towards a later timing after exposure to persistent delays (*i.e.*, 250 ms) between the pulling action of one hand and the grip force of the other. Given that previous experiments demonstrated that this grip modulation is anticipatory and time-locked to the predicted self-generated change in the load force by the internal model ^41–45^, these results suggested that the internal model is recalibrated by learning the delay between the pulling action and the grip response ^40^. In strong agreement, Kilteni et al. (2019) showed that nondelayed self-generated touches produced by touching one hand with the other feel stronger and are more frequently reported as ticklish after exposure to persistent delays (*i.e.*, 100 ms) between the action of one hand and resulting touch on the other. Without exposure to delays, nondelayed self-generated touches are robustly attenuated compared to externally generated touches ^17,46–66^ or delayed self-generated touches of identical intensity ^46,59,61,65,67,68^ because the nondelayed self-generated touches are received at the predicted time of contact between the two hands by the internal model. Consequently, the increased magnitude of the nondelayed self-generated touches after exposure to persistent delays suggested that the predicted time of contact shifted ^46^. In further support of this interpretation, Kilteni et al. (2019) showed that during the same exposure to the delays, delayed self-generated touches become expected and thus attenuated, resulting in similar perceptual responses between delayed and nondelayed touches. Together, these studies ^40,46^ provided evidence that persistent sensorimotor delays can be learnt to be expected, forcing a recalibration in the predictions of the internal model.

Nevertheless, how the brain learns to adapt to these delays remains unclear as, to our knowledge, no neuroimaging study has assessed adaptation-related changes during delays in self-generated touches. Instead, previous neuroimaging studies showed that when brief delays were introduced between participants’ movement and the self-generated touches, there were increased cerebellar ^67^ and somatosensory ^65^ responses, increased functional connectivity between the supplementary motor area and cerebellum, and decreased functional connectivity between the somatosensory cortices and cerebellum ^65^. Critically, both studies were designed to specifically preclude adaptation to the delay, either by alternating trials of different delays in random order ^67^ or by blocking together delayed trials ^65^ that were purposefully selected to be too few to trigger adaptation ^46^.

Here, we used our previous sensorimotor self-touch delay adaptation paradigm ^46^ together with functional magnetic neuroimaging (fMRI) to examine the effect of systematic exposure to delays on the neural responses associated with self-touch. We analyzed BOLD responses evoked when participants administered with their right index finger nondelayed or delayed (*i.e.*, 100 ms) self-generated touches to their left index finger (through a setup, see below). Critically, nondelayed and delayed self-generated touches occurred in two contexts, either while participants were being systematically exposed to no injected delay (0 ms delay, *baseline* session) or when they were systematically exposed to persistent delays of 100 ms (*adaptation* session) between the movement of the right index finger and the resulting touch on the left index finger.

By adopting this experimental design, we could test three specific hypotheses. First, we hypothesized that if participants update their sensorimotor predictions during the *adaptation* session, delayed self-generated touches should become neurally attenuated, while nondelayed touches should exhibit less neural attenuation. These changes should be reflected in dynamic alterations in neural activity within the somatosensory cortices, resulting in more comparable somatosensory activity levels for the two types of touches after the delay exposure. Second, given the involvement of the cerebellum in acquiring, storing, and recalibrating the internal models, as well as in sensorimotor learning ^16,25,69–76^, we expected delay adaptation to produce plastic changes in cerebellar activity reflecting the temporal recalibration of the internal model. Third, we reasoned that once the internal model has been recalibrated to a certain extent and the participants’ nondelayed self-touches are “treated” as more unexpected, these will elicit activity in areas involved in conflict monitoring, such as the anterior cingulate cortex ^33,77–83^. This would occur because the unexpected nondelayed self-touch is conflicting with a lifetime of experiencing (nondelayed) self-touches as fully predictable.

## Materials and Methods

### Participants

We aimed for a sample size of 30 healthy participants based on our previous study ^50^. However, due to a major scanner failure, one participant was not scanned at all. Five participants were further excluded: one for exiting the scanner in the middle of the fMRI runs, one for reporting extreme sleepiness during some of the fMRI runs, and three for technical problems with the setup or with registering their responses. Consequently, the behavioral and fMRI analyses included data from a total of 24 participants (12 women, 12 men; 22 right-handed, 2 ambidextrous; 19-36 years old). The Ethics Review Authority approved the study (project: #2016/445-31/2, amendment: #2018:1397-32). Handedness was assessed using the Edinburgh Handedness Inventory ^84^, and all participants provided their written informed consent.

### Experimental design

Throughout all fMRI and behavioral runs, participants lay comfortably on the MRI scanner bed with their left hands placed palm-up on an MR-compatible plastic table and their left index finger in contact with a 3D-printed probe that contained a force sensor (**Figure 1a**). The probe was controlled by an electric motor through string-based transmission. The participants’ right hand was resting palm down on a support directly above the left hand; the right index finger was placed next to a second force sensor (identical sensor to the sensor used for the left index) placed on the table on top of (but not in contact with) the probe on the left index finger (10 cm approximately). Both arms were supported by sponges to maximize the comfort of the participants. The participants’ right arm and hand were peripherally visible. To prevent head motion, the participants’ head was fixed with foam pads placed between their ears and the head coil. All participants wore earplugs and a pair of headphones to protect their hearing from the scanner noise.

**Figure 1.**
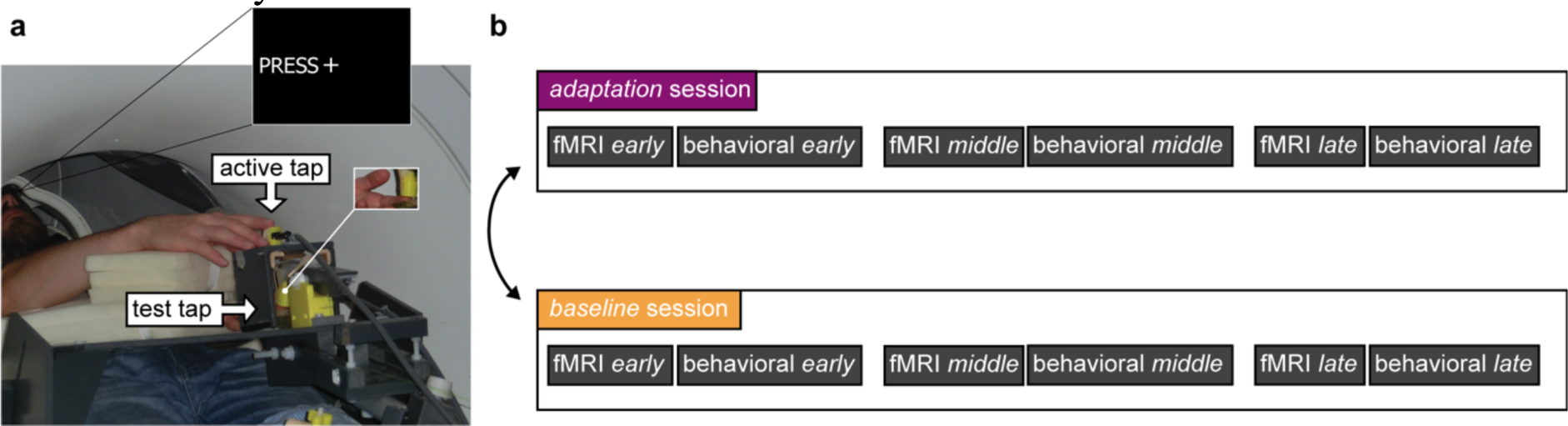
Experimental setup and design. (a) An fMRI-compatible setup was used in all the fMRI and behavioral runs. In all runs, participants were asked to tap a force sensor with their right index finger (active tap) that triggered the electric motor to apply a tap on their left index finger (test tap; magnified view outlined in small white box). **(b)** The study was organized in two sessions, each consisting of 3 fMRI and 3 behavioral runs (*early*, *middle*, *late*). In the *adaptation* session (purple), we injected a 100 ms delay between the active tap and the test tap, while in the *baseline* session (orange), we injected a 0 ms delay.

In all fMRI and behavioral trials, participants tapped a force sensor with their right index finger (active tap) to trigger a force on the left index finger (test tap) (**Figure 1a**). There were two sessions: the *adaptation* session that involved continuous exposure to an injected delay of 100 ms between the active tap and the test tap (purple, **Figure 1b**) and the *baseline* session that involved continuous exposure to a 0 ms injected delay (no injected delay) between the active tap and the test tap (orange, **Figure 1b)** and served as the control session. The order of the two sessions was randomized across participants. In the final sample, 14 participants started with the *baseline* session and 10 participants started with the *adaptation* session.

Each session consisted of three fMRI runs (*early*, *middle*, *late*) and three behavioral runs (*early*, *middle*, *late*) in alternating order, resulting to a total of 6 fMRI and 6 behavioral runs per participant (**Figure 1b**). We deliberately chose to have separate behavioral and fMRI runs in order to reduce movement-related artifacts (*e.g.*, head motion) associated with verbal responses in the behavioral task and to ensure that the BOLD signal reflected activation related to the basic sensorimotor task of self-touch and the processing of somatosensory signals and not high-level cognitive processes engaged in decision-making in the force-discrimination task (*e.g.*, working memory, decision making) (see Behavioral runs). Within each session, all three fMRI runs were identical, and all three behavioral runs were identical. Before the 6 fMRI and behavioral runs reported here, additional fMRI and behavioral runs were conducted as part of a different study ^65^.

### System delay and experimental delays

There was an intrinsic delay of 53 ms ^65^ in our fMRI-compatible setup. This means that both sessions included a 53 ms intrinsic delay but differed in whether we injected a 0 ms or 100 ms delay. Note that delays of ∼ 50 ms have been previously shown to not impact somatosensory attenuation compared to smaller delays (*e.g.*, 11 ms) ^59^, suggesting that our 0 ms injected delay session simulated natural self-touch well ^65^.

The size of the injected delay (*i.e.*, 100 ms) in the *adaptation* session was chosen for three reasons. First, previous studies showed that delays of 100 ms are long enough to perturb somatosensory attenuation ^46,59,61,65^. Second, delays on the order of 100 to 200 ms are typically unnoticed by participants ^61^; this enables the experimental manipulation to be covert, prevents higher-level cognitive processes associated with detecting longer delays from potentially interfering with the basic sensorimotor temporal recalibration ^38^, and avoids evoking fMRI activations in areas related to exogenous attention due to unexpected perceptual changes. Third, and more importantly, we previously showed that participants can adapt to a 100 ms injected delay after repeated exposure. This adaptation was reflected in two correlated findings: the perceived magnitude of the otherwise nonattenuated delayed self-generated touch (100 ms) was attenuated after delay adaptation and the perceived magnitude of the otherwise attenuated nondelayed self-generated touch (0 ms) was less attenuated after delay adaptation ^46^.

### Procedures

#### fMRI runs

As mentioned earlier, each session had three identical fMRI runs – the *early*, *middle,* and *late* runs – each consisting of 235 trials, on average, and including trials of both nondelayed and delayed self-generated touches. In each trial, participants tapped the force sensor with their right index finger (active tap) after an auditory GO cue. A “PRESS” message was displayed on the screen seen through a mirror attached to the head coil to remind participants’ what they had to do after the auditory cue (**Figure 2**). The active tap of the right index finger (force exceeding > 0.4 N) triggered the test tap with either a 100 ms injected delay (**Figure 2**, delayed) or with a 0 ms injected delay simulating self-touch (**Figure 2**, nondelayed). Prior to the experiment, we asked participants to tap the sensor with their right index finger at an intensity that is comfortable to maintain throughout all runs. The test taps had a fixed intensity of 2 N and were administered for 250 ms (mean ± s.e.m.: 246.694 ± 4.755 ms).

**Figure 2.**
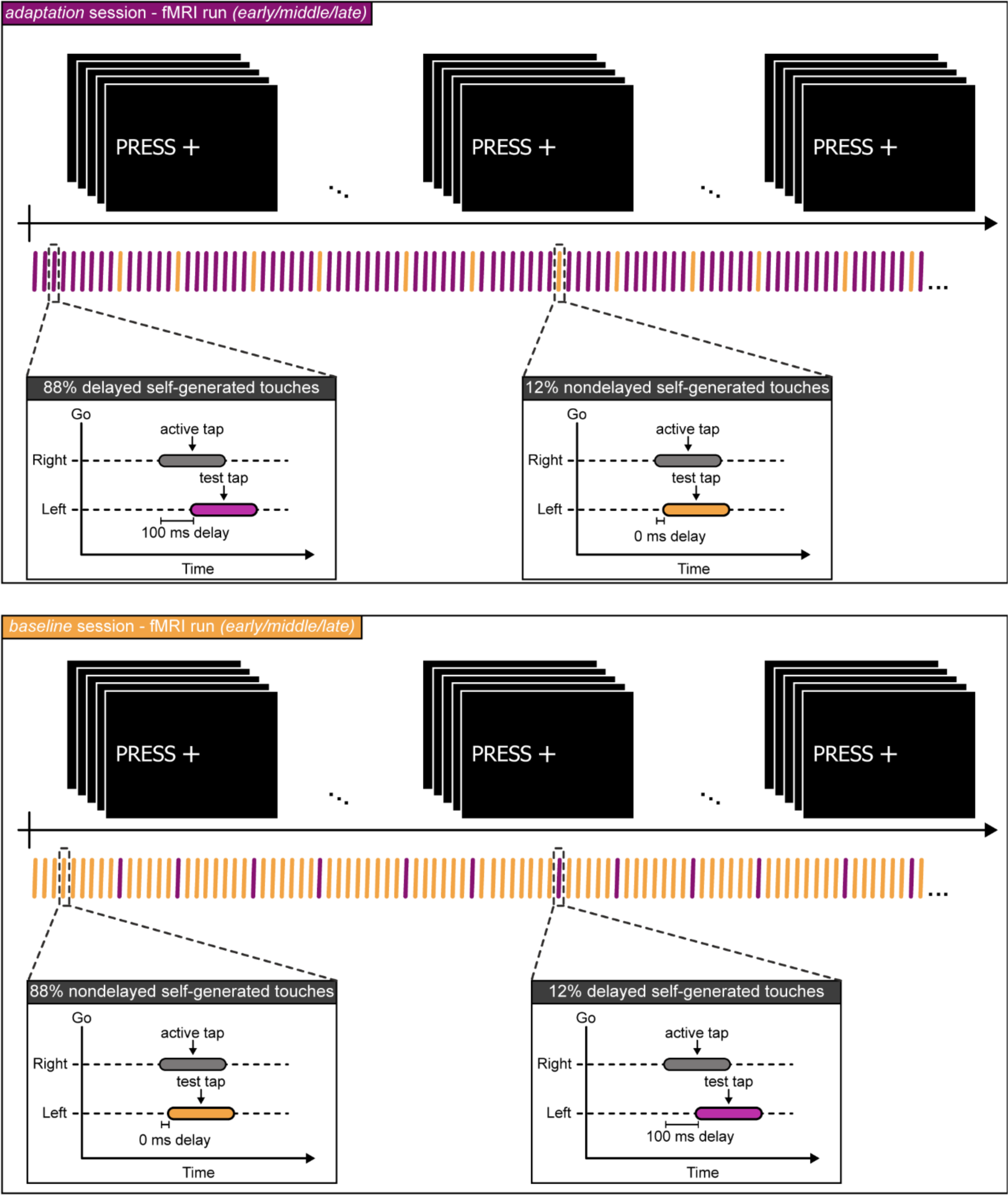
The fMRI runs. Each session consisted of three identical fMRI runs **–** the *early*, *middle*, and *late* fMRI runs. Each run consisted of a series of delayed (purple) and nondelayed (orange) self-generated touches. Participants were asked to fixate their gaze on the fixation cross seen on the screen. In every trial, participants heard an auditory GO cue and received the message “PRESS” on the screen that instructed them to tap the force sensor with their right hand (active tap, gray rectangle) and received the test tap on their left index finger (2 N) either with a 0 ms (orange rectangle) or a 100 ms injected delay (purple rectangle) from the electric motor. Each run included the same percentage of nondelayed and delayed self-generated touch trials. In the *adaptation* session (purple, top), 88% of trials were delayed (purple) and 12% were nondelayed (orange) self-generated touches. In the *baseline* (orange, bottom), 88% of the trials were nondelayed self-generated touches and 12% of the trials were delayed self-generated touches.

In the *adaptation* session, 88% of the trials were delayed self-generated touches (*i.e.*, we injected a 100 ms delay between active and test taps), while 12% of the trials were nondelayed self-generated touches (*i.e.*, we injected a 0 ms delay between active and test taps). Each nondelayed self-generated touch pseudorandomly appeared after 5 to 9 delayed self-generated touches. By contrast, in the *baseline* session, 88% of the trials were nondelayed self-generated touches and 12% of the trials were delayed self-generated touches. Here, each delayed self-generated touch pseudorandomly appeared after 5 to 9 nondelayed self-generated touches. This design thus provided systematic exposure to the delay while enabling us to assess neural responses to both delayed and nondelayed self-generated touches in each session.

#### fMRI data acquisition

fMRI acquisition was performed using a General Electric 3T scanner (GE750 3T) equipped with an 8-channel head coil. Gradient echo T2*-weighted EPI sequences with BOLD contrast were used as an index of brain activity. A functional image volume was composed of 42 slices (repetition time: 2000 ms; echo time: 30 ms; flip angle: 80°; slice thickness: 3 mm; slice spacing: 3.5 mm; matrix size: 76 × 76; in-plane voxel resolution: 3 mm).

A total of 155 functional volumes were collected for each participant during each run, resulting in a total of 930 functional volumes (155 volumes × 6 runs). For the anatomical localization of activations, a high-resolution structural image containing 180 slices was acquired for each participant before the acquisition of the functional volumes (repetition time: 6.404 ms; echo time: 2.808 ms; flip angle: 12°; slice thickness: 1 mm; slice spacing: 1 mm; matrix size: 256 × 256; voxel size: 1 mm × 1 mm × 1 mm).

#### Behavioral runs

As with the fMRI runs, each session included three identical behavioral runs – the *early*, *middle,* and *late* runs (**Figure 3**). Each behavioral run included trials of nondelayed and delayed self-generated touches and 50 response trials. The trials of nondelayed and delayed self-generated touches were identical to those of the fMRI runs and served to retain the delay adaptation and avoid washout effects - in line with a neural model of temporal sensorimotor recalibration ^34^ and our previous study ^46^. In the three behavioral runs of the *adaptation* session, all trials were delayed self-generated touches, while in the three behavioral runs of the *baseline* session, all trials were nondelayed self-generated touches.

**Figure 3.**
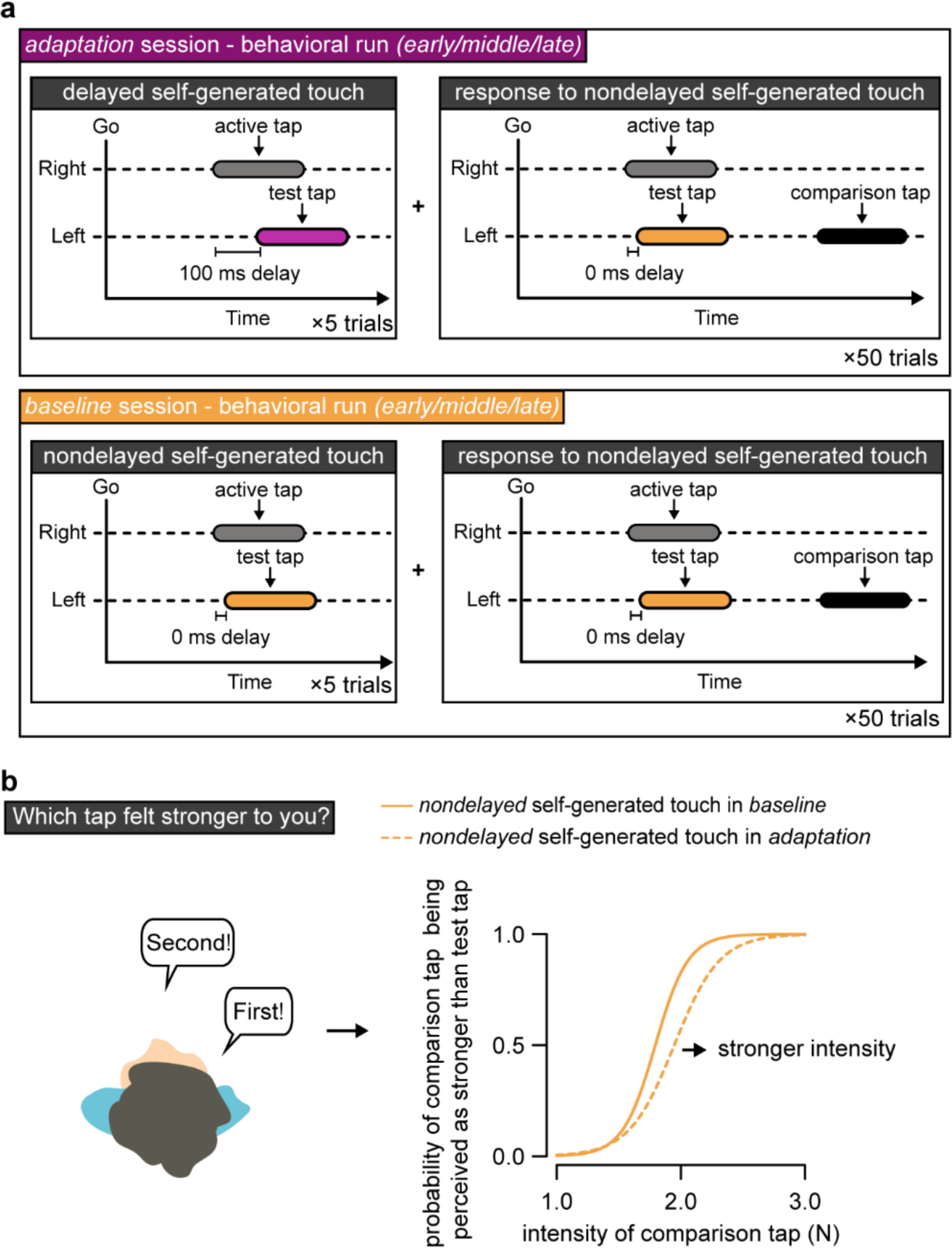
The behavioral runs. (a) Each behavioral run consisted of 50 response trials with each response trial being preceded by 5 delayed self-generated touch trials in the *adaptation* session (top, purple) or 5 nondelayed self-generated touch trials in the *baseline* session (bottom, orange). Identical to the fMRI trials, upon hearing an auditory GO cue, participants tapped the force sensor with their right index finger (active tap) and received the test tap on their left index finger (2 N) with delay (*adaptation* session) or without delay (*baseline*). In the response trials, the test tap was always nondelayed (orange). Following, a second tap (comparison tap) of variable magnitude (black rectangle) was applied to the participant’s left index finger. **(b)** After receiving both taps, participants verbally reported which of the two taps (*i.e.*, the test or the comparison tap) felt stronger. Therefore, the behavioral runs enabled us to track changes in the perception of nondelayed self-generated touch due to delay adaptation. **(c)**The participants’ responses on each task were fitted with a logistic function for each run of the baseline (solid line) and adaptation session (dashed line). The orange color of the two lines represents that they both concern the perception of nondelayed self-generated touch.

The 50 response trials enabled us to collect perceptual responses and included a two-alternative forced-choice force-discrimination task (**Figure 3**) that has previously been used to quantify somatosensory perception ^46,51,53–55,58,59,65,66,68^, including during delay adaptation ^46^. Each response trial was preceded by five delayed (**Figure 3a**, top, purple) or nondelayed (**Figure 3b**, bottom, orange) self-generated touches depending on the session. In the response trials, participants were presented with two taps on the left index finger – the nondelayed self-generated test tap of fixed magnitude (2 N) and one externally generated comparison tap of variable magnitude (1, 1.5, 1.75, 2, 2.25, 2.5, or 3 N). After receiving the two taps, participants verbally indicated which tap felt stronger (**Figure 3b** left). A “JUDGE” message was displayed on the screen seen through a mirror attached to the head coil to remind participants’ that they had to report their judgments. As in the fMRI runs, the taps were administered for 250 ms (mean ± s.e.m.: 251.944 ± 4.071 ms). Within the 50 response trials of each behavioral run task, each level of comparison tap (1, 1.5, 1.75, 2, 2.25, 2.5, or 3 N) was repeated 7 times, except for the level of 2 N, which was repeated 8 times.

#### Post-experiment debriefing

At the end of the experiment, once all fMRI and behavioral runs were concluded, participants were asked whether they perceived that we injected a delay between the active tap of the right index finger and the test tap on the left index finger in any of the runs. Only one participant reported perceiving the presence of the injected delays, similar to previous studies reporting limited awareness for such brief delays in self-generated touches or tones ^61,85^.

### Data analysis

#### fMRI data preprocessing

A standard preprocessing pipeline was used, including realignment, unwarping and slice-time correction using Statistical Parametric Mapping 12 (SPM12; Welcome Department of Cognitive Neurology, London, UK, http://www.fil.ion.ucl.ac.uk/spm) software. Outlier volumes were detected using the Artifact Detection Tools, employing the option for liberal thresholds (global-signal threshold of *z* = 9 and subject-motion threshold of 2 mm). Next, we simultaneously segmented the images into gray matter, white matter and cerebrospinal fluid and normalized them into standard MNI space (Montreal Neurological Institute, Canada). Subsequently, the images were spatially smoothed using an 8-mm FWHM Gaussian kernel. The structural images were also simultaneously segmented (into gray and white matter and cerebrospinal fluid) and normalized to MNI space.

#### Univariate data analysis and hypotheses

BOLD signal responses were modeled by fitting voxelwise GLMs to the data of each fMRI run. For all six runs, the main regressor of interest was deviant touches (*i.e.*, 12% delayed self-generated touches in the *early*, *middle*, and *late* runs of the *baseline* session, and 12% nondelayed self-generated touches in the *early*, *middle*, and *late* runs of the *adaptation* session), while the repeated standard touches (*i.e.*, 88% nondelayed self-generated touches in the *early*, *middle*, and *late* runs of the *baseline* session, and 88% delayed self-generated touches in the *early*, *middle*, and *late* runs of the *adaptation* session) were modelled as the “implicit” baseline. The onset of the trials was defined as the time when the magnitude of the *test* tap peaked, their duration was set to zero, and they were convolved with the canonical hemodynamic response function of SPM12. Any trials in which the participants did not tap the sensor with their right index finger after the auditory cue, tapped too lightly to trigger the touch on the left index finger (active tap < 0.4 N), tapped more than once, or tapped before the auditory GO cue were excluded from the regressor of interest and implicit baseline and modeled as four (4) individual regressors of no interest. This resulted in the exclusion of 805 trials out of 33877 fMRI trials (2.4%) from the main regressor of interest and the implicit baseline. In addition, the six motion parameters and any outlier volumes were included as regressors of no interest. To account for the potential influence of small variations in the magnitude of the self-generated force of active taps on the BOLD signal ^86,87^, we also included the magnitude of the active tap on each trial as a parametric modulator. Finally, the first-level analysis was restricted to gray matter voxels using a binary (threshold of 0.2) and smoothed mask (8-mm FWHM Gaussian kernel) of gray matter, which was based on the individual’s segmented structural image (gray matter).

Our overall hypothesis was that systematic exposure to sensorimotor delay in the *adaptation* session would force the internal model to recalibrate its temporal predictions about the somatosensory feedback towards the delayed timing. This recalibration would render the nondelayed self-touches as unpredicted touches that are subject to less attenuation and whose presence elicits conflict-related activity instead. More specifically, we expected that as participants adapted to the delay, somatosensory responses to the nondelayed touches would increase (*i.e.*, less attenuation) and responses to the delayed touches would decrease (*i.e.*, attenuation), resulting in comparable responses in the right somatosensory cortices for both types of self-generated touches. In accordance, behavioral responses to the nondelayed touches would also be stronger (*i.e.*, less attenuation). Second, we anticipated that the recalibration of the internal model would be associated with adaptation-related changes in cerebellar responses to nondelayed self-generated touches in the *adaptation* session. These changes were specifically expected (a) in lobules I-VI and VIII given that these lobules are considered to contain two separate motor representations of the body ^88–96^ and (b) in the left hemisphere because they concern the recalibration of touches applied on the left hand and the cerebellum contains ipsilateral body representations ^88,97^. Third, we theorized an increase in the activity of the anterior cingulate cortex (ACC) towards the end of the *adaptation* session, showing that the otherwise natural nondelayed self-touches are now eliciting conflict-related activity because of the internal model recalibration ^33^. Based on these hypotheses, we formulated planned neuroanatomically directed hypotheses (defined ROIs for small volume correction, see further below), in addition to a whole-brain explorative approach.

With respect to the time course of activity changes, in our previous behavioral study ^46^, we showed adaptation effects for delayed self-generated touches after 750 exposure trials (50 initial exposure trials + 2 × 5 × 70 re-exposure trials), and for nondelayed self-generated after 1250 exposure trials (200 initial exposure trials + 3 × 5 ×70 re-exposure trials). Assuming a similar time-course, we expected signs of delay adaptation in the *late* run (> 914 trials = 207 per fMRI run × 2 runs + 250 trials per behavioral run × 2 runs) but not in the *early* run (∼ 207 trials) or *middle* run (∼ 664 trials). In other words, the adaptation-related somatosensory changes should be present in the *late* run, the recalibration of the internal model (*i.e.*, left cerebellum) should occur earlier than the *late* run (*e.g.*, *early* or *middle* run) and error-related activity (*i.e.*, ACC) in response to nondelayed self-touches should peak at the end of the adaptation (*i.e.*, *late* run).

Six contrasts against the implicit baseline were created for all the deviant self-generated touches: *baseline_early_* > 0, *baseline_middle_* > 0, *baseline_late_* > 0, *adaptation_early_* > 0, *adaptation_middle_* > 0, *adaptation_late_* > 0. These contrasts were inserted in a 2 × 3 full factorial design with two within-subjects factors: session (*adaptation* or *baseline*) and run (*early*, *middle*, *late*). Based on our previous results ^65^, we first confirmed that the delayed self-generated touches elicit stronger somatosensory activity than the nondelayed ones in the *baseline* session (*i.e.*, *baseline* > 0, *baseline_early_* > 0, *baseline_middle_* > 0, *baseline_late_* > 0), that is, when participants were not systematically exposed to the delayed touches but to the nondelayed ones. Next, we checked whether this difference existed in the *adaptation* session as well or whether delayed and nondelayed self-generated touches elicited comparable responses in the right somatosensory cortices (*i.e.*, *adaptation* < 0, *adaptation_early_* < 0, *adaptation_middle_* < 0, *adaptation_late_* < 0). To provide support for comparable BOLD responses to delayed and nondelayed touches, we extracted the average activity from somatosensory areas revealed in the *baseline* session using the Marsbar Toolbox ^98^, and we performed one-tailed Bayesian comparisons (*BF_0-_*, default Cauchy prior of 0.707) for the difference in the *adaptation* session being smaller than zero (*i.e.*, BOLD responses will be greater for delayed than nondelayed touches, as in the SPM contrast). We further directly contrasted the same difference in somatosensory responses (delayed - nondelayed touches) between the *baseline* and the *adaptation* sessions. Finally, to specifically test for effects related to the length of the exposure, we created and tested the contrasts between the runs of each session (*i.e.*, *baseline_late_* > *baseline_middle_*, *baseline_late_* > *baseline_early_*, *baseline_middle_* > *baseline_early_*, *adaptation_late_* > *adaptation_middle_*, *adaptation_late_* > *adaptation_early_*, *adaptation_middle_* > *adaptation_early_*). For transparency, we also report the same contrasts in the opposite direction.

We performed small-volume corrections within regions of interest (ROIs). The somatosensory ROIs included the right primary somatosensory cortex, defined as a spherical region of 10-mm radius, centered at a peak detected in our previous study (MNI coordinates: *x* = 50, *y* = -20, *z* = 60) using the same scanner, same equipment, and same tactile stimulation (2 N) applied to the same finger (left index finger) ^50^. The right secondary somatosensory cortex was defined using the Anatomy Toolbox ^99^ by selecting the Brodmann area OP1(SII). Two ROIs included the hemispheres of the left cerebellar lobules IV-VI and VIII, and one ROI included the anterior cingulate cortex, all defined with the Wake Forest University Pickatlas toolbox ^100^.

In addition to these ROIs, we also report analyses at the whole-brain level. For each peak activation, the coordinates in MNI space, the *z* value and the *p* value are reported. We denote that a peak survived a threshold of *p* < 0.05 after correction for multiple comparisons at the whole-brain or small-volume level by adding the term “FWE-corrected” after the *p* value.

#### Performance during the scans

To assess whether the participants pressed with forces of similar magnitude during the six fMRI and the six psychophysical runs, and thus rule out that any perceptual or neural effects are driven by differences in the produced or received forces rather than the injected delay, for each trial, we extracted the peak amplitudes of the test and active taps (defined as the peak force recording of each force sensor within the trial).

#### Labelling for anatomical areas

Macroanatomical labels for anatomical reference of significant activations were made using the SPM Anatomy Toolbox ^99^. The anatomical locations of the activation peaks were also directly compared with a mean structural MRI scan of this group of participants’ brains to verify correct anatomical labelling with respect to major sulci and gyri in this group.

#### Behavioral data analysis and hypotheses

There were no missing trials in any of the six psychophysical tasks, resulting in a total of 7200 trials (24 subjects × 50 response trials × 6 tasks = 7200 trials). After data collection, we excluded any psychophysical trials in which the participants did not tap the sensor with their right index finger after the GO cue, tapped too lightly to trigger the touch on the left index finger (active tap < 0.4 N), tapped more than once or tapped before the GO cue, as well as any trials in which the test tap was not applied correctly (test tap <1.85 N or test tap > 2.15 N). This resulted in the exclusion of 380 of 7200 psychophysical trials (5.27%).

We fitted the participants’ responses with a generalized linear model using a *logit* link function (Equation 1) (**Figure 1d**, *right*):

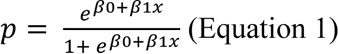

Before fitting the responses, the *comparison* forces were binned to the nearest of the seven possible comparison tap intensities (1, 1.5, 1.75, 2, 2.25, 2.5, or 3 N). We extracted two parameters of interest: the point of subjective equality 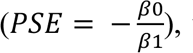, which represents the intensity at which the *test* tap felt as strong as the *comparison* tap (*p* = 0.5) and quantifies the perceived intensity of the *test* tap, and the just noticeable difference 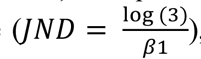, which reflects the participants’ discrimination capacity. PSEs and JNDs are independent sensory judgments: higher PSE values indicate a stronger perceived magnitude, while higher JND values indicate a lower force discrimination capacity (*i.e.*, lower somatosensory precision).

We used a repeated measures ANOVA (RM ANOVA) to analyze our data with two repeated factors: the session (*adaptation* or *baseline*) and the run (*early*, *middle*, *late*). Based on our previous study ^46^, we expected to find a significant increase in the perceived magnitude of nondelayed self-generated touch (PSE) in the *adaptation* compared to the *baseline* session (main effect of session). In addition, in our previous study, we found the earliest behavioral effects on the perception of *nondelayed* self-generated touch after a total of 1250 exposure trials (200 initial exposure trials + 3 × 350 re-exposure trials = 1250 total exposure trials) (Experiment 2 in ^46^). Consequently, as in the fMRI runs, we here expected to see a significant increase in the perceived magnitude of nondelayed self-generated touch in the *late* run that was performed after > 1100 trials of exposure to the delayed touches, but not earlier. Finally, we did not expect to find any effects for any other variables: JNDs or active taps’ or test taps’ magnitude.

In addition to the RM ANOVA, planned pairwise comparisons were conducted using parametric (paired *t-*tests) or nonparametric analyses (Wilcoxon sign rank tests) depending on the data normality, which was assessed using the Shapiro–Wilk test. For each test, 95% confidence intervals (*CI*^95^) are reported. Bayesian RM ANOVAs and t tests were used to assess evidence for the null hypothesis compared to the alternative hypothesis (*BF_01_*).

For all Bayes’ factors reported for the fMRI and behavioral runs, we interpret a factor 1 to 3 as ‘anecdotal’, 3 to 10 as ‘moderate’, 10 to 30 as ‘strong’, and greater than 100 as ‘decisive’, indicating anecdotal, strong, or decisive support for the null hypothesis, respectively ^101^.

### Software

fMRI data preprocessing and analysis was performed using SPM12 on MATLAB R2019b. Python 3.11.3 (libraries NumPy, SciPy) was used to preprocess the behavioral data. R (version 4.2.2, 2022-10-31) ^102^ and JASP 0.16.4 ^103^ were used for statistical analysis of the behavioral data. The flat representation of the human cerebellum (cerebellar flatmap) provided with the SUIT toolbox was used to visualize the group average imaging data ^104^.

## Results

### Comparable force production and force stimuli across all behavioral and fMRI runs

We first confirmed that participants received comparable touches (test taps) and performed comparable presses (active taps) across all behavioral and fMRI runs (**Supplementary Figures S1, S2).** This was supported by both frequentist and Bayesian analyses (**Supplementary Texts S1, S2**), ensuring that any perceptual or neural effects detected during the runs were not driven by the physical magnitude of the touch participants received on their left index finger (test taps) or by the magnitude of the participants’ presses of their right index finger (active taps).

### When systematically exposed to delayed self-generated touches, nondelayed self-generated touches feel stronger over time compared to the same touches in the baseline

If participants adapt to the sensorimotor delay and predict their somatosensory feedback to arrive at the delayed time, then the nondelayed self-generated touches should be subject to less attenuation because they are received at a timing that is now unexpected and thus perceived as stronger over the exposure time. Confirming our hypothesis, a two-way RM ANOVA on the PSEs with the session and run as repeated factors revealed a significant main effect of session (*F*(1, 23) = 4.863, *p* = 0.038); that is, nondelayed self-generated touches felt significantly stronger in the *adaptation* session compared to identical nondelayed self-generated touches in the *baseline*, replicating our previous effects ^46^ (**Figure 4a, Supplementary Figure S3**). The main effect of run was not significant (*F*(2, 46) = 1.810, *p* = 0.175), suggesting that time *per se* was not driving any perceptual affects. In agreement with our hypothesis and previous results ^46^, the difference between the PSEs of the two sessions was more prominent in the *late* runs (paired t-test, *n* = 24, *t*(23) = 3.468, *p* = 0.002, *CI*^95^ = [0.047, 0.185]) than in the *early* (paired t-test, *n* = 24, *t*(23) = 1.045, *p* = 0.307, *CI*^95^ = [-0.038, 0.114]) or *middle* runs (paired t-test, *n* = 24, *t*(23) = 0.659, *p* = 0.51, *CI^95^* = [-0.055, 0.107]) (**Figure 4b**), although we note that the interaction session × run did not reach statistical significance (*F*(2, 46) = 2.695, *p* = 0.078, *BF_01_* = 1.295). The change in the PSEs between the two sessions was time-dependent, as it was not apparent between the *early* and the *middle* trials (*paired t-test*, *n* = 24, *t*(23) = -0.317, *p* = 0.754, *CI^95^* = [-0.095, 0.069]), it became numerically greater in the *late* compared to the *early* runs (*paired t-test*, *n* = 24, *t*(23) = 1.738, *p* = 0.096, *CI^95^* = [-0.015, 0.170]), and significantly greater in the *late* compared to *middle* runs (*paired t-test*, *n* = 24, *t*(23) = 2.156, *p* = 0.042, *CI^95^*= [0.004, 0.117]) (**Figure 4c**). As expected, the delay adaptation was specific to the perceived magnitude (PSEs) of the self-generated touch and did not affect its discrimination capacity (JNDs): there was no statistically significant main effects of session (*F*(1, 23) = 2.472, *p* = 0.130) or run (*F*(2, 46) = 1.047, *p* = 0.359), nor a significant interaction effect (*F*(2, 46) = 0.121, *p* = 0.886) (**Figure 4d**), strongly supported by a Bayesian two-way RM ANOVA (*BF_01_* = 29.012). **Figure 4e** illustrates the abovementioned results as the shift in the group psychometric curve observed in the *late* phase of the *adaptation* session.

**Figure 4.**
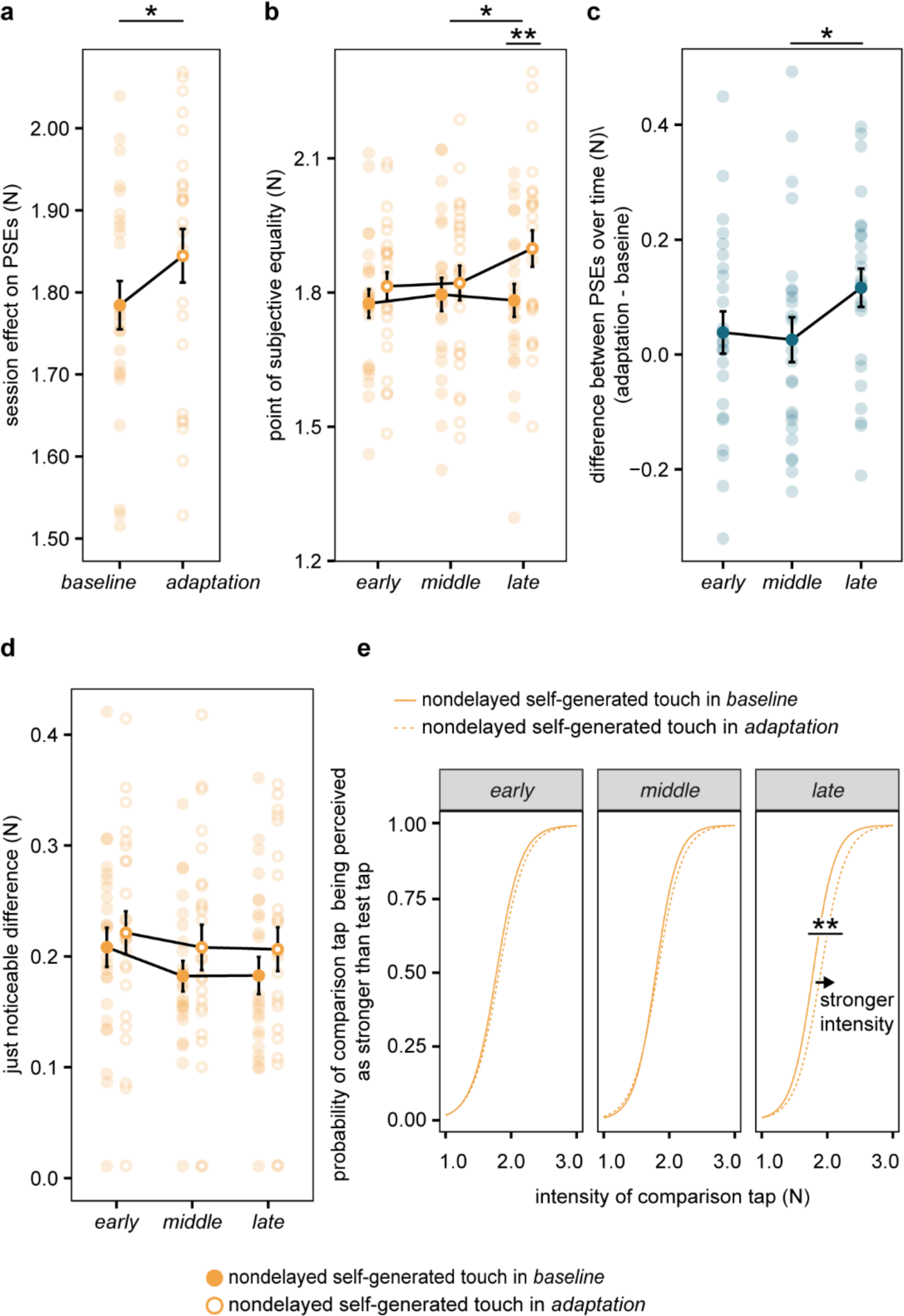
Perceptual changes to nondelayed self-generated touches due to exposure to delayed self-generated touches. (a) Individual and group PSE values (mean ± s.e.m.) averaged across all runs, showing a significant increase (*p* = 0.038) in the perceived magnitude of the nondelayed self-generated touch after repeated exposure to the 100 ms injected delay (empty circles) compared to the *baseline* session (filled circles). **(b)** Individual and group PSE values (mean ± s.e.m.) per session and run showing that the nondelayed self-generated touches felt significantly stronger in the *late* runs of the *adaptation* session compared to the *baseline* (*p* = 0.002). **(c)** Individual and group PSE differences between the two sessions (mean ± s.e.m.) over time. The change was significantly greater in the *late* compared to *middle* runs (*p* = 0.042). **(d)** Individual and group JND values (mean ± s.e.m.) per session and run depicting no statistically significant changes in JNDs in any of the runs or sessions. **(e)** Group psychometric curves per session and run based on the group PSE and JND values. The arrow indicates the direction of the shift in the perceived magnitude of the nondelayed self-generated touch during the *late* runs of the *adaptation* session (dashed line) compared to the *baseline* session (solid line). (* *p* < 0.5, ** *p* < 0.01)

In conclusion, the nondelayed self-generated touches were significantly less attenuated during the *late* phase of the *adaptation* compared to identical self-generated touches in the *baseline* session, effectively demonstrating behavioral adaptation to the systematic sensorimotor delay that was scaled with longer exposure. This confirms that we could replicate our previous behavioral results ^46^ in the current setup adopted for the MRI-scanner environment.

### Responses in contralateral secondary somatosensory cortex are stronger for delayed than nondelayed self-generated touches in the baseline and do not change over time

When participants are not systematically exposed to delayed self-generated touches but to nondelayed ones (*baseline* session), there should be no delay adaptation, as there is no sensorimotor delay to adapt to. Consequently, somatosensory BOLD responses to delayed self-generated touches should be stronger compared to nondelayed self-generated touches ^65^ and without changes over time. Indeed, somatosensory responses in the right secondary somatosensory cortex were significantly stronger for delayed compared to nondelayed self-generated touches (*baseline* > 0, MNI: *x* = 54, *y* = -20, *z* = 22, *p* < 0.001 FWE-corrected at the whole brain level) (**Figure 5a-b, Supplementary Table S1**). A similar pattern was observed in the responses of the left secondary somatosensory cortex (MNI: *x* = -44, *y* = -30, *z* = 20, *p* < 0.001 uncorrected) and the right S1 (MNI: *x* = 54, *y* = -16, *z* = 44, *p* = 0.024 uncorrected) that did not survive whole-brain corrections. The effects in right secondary somatosensory cortex were statistically observed (*p* < 0.05 FWE-corrected) in all runs: *early* (*baseline_early_* > 0, MNI: *x* = 58, *y* = -16, *z* = 20, *p* = 0.004 FWE-corrected), *middle* (*baseline_middle_* > 0, MNI: *x* = 44, *y* = -28, *z* = 22, *p* = 0.006 FWE-corrected), and *late* (*baseline_late_* > 0, MNI: *x* = 54, *y* = -20, *z* = 22, *p* < 0.001 FWE-corrected). Critically, when testing for any differences in the BOLD responses over the three runs of the *baseline* session, there were no significant changes from the *early* to the *middle* run (**Supplementary Table S2)**, from the *middle* to the *late* run (**Supplementary Table S3)**, nor from the *early* to the *late* run (*i.e.*, no activations at *p* > 0.001 uncorrected). There were also no significant activations in the opposite direction: *i.e.*, from *late* to *middle*, *late* to *early* and *middle* to *early* (*i.e.*, no activations at *p* < 0.001 uncorrected for the three contrasts).

**Figure 5.**
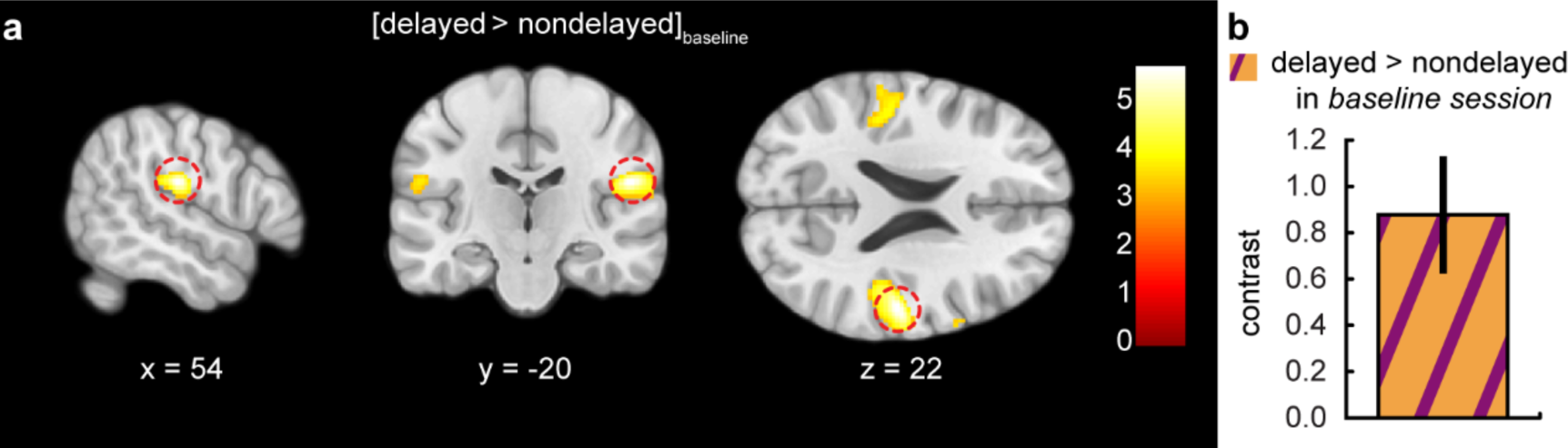
Somatosensory activations during the delayed compared to the nondelayed self-generated touches in the *baseline* session. (a) Sagittal, coronal, and axial views of the significant peak of activation (*p* < 0.05 FWE-corrected) located in the right secondary somatosensory cortex (parietal operculum; MNI: *x* = 54, *y* = -20, *z* = 22) during all trials of the *baseline* session. The activation maps are rendered on the MNI-152 template brain and are displayed at an uncorrected threshold of *p* < 0.001. Red circles are centered over the main significant peak. The color bar indicates the values of the t-statistic. **(b)** Contrast estimates and 90% CI for the significant somatosensory peak in all runs of the *baseline* session.

Together, these results demonstrate that without systematic exposure to delayed self-generated touches (*baseline* session), somatosensory responses in the right secondary somatosensory cortex are stronger for delayed than for identical nondelayed self-generated touches and remain stable over time. This replicates the well-established somatosensory attenuation effect to self-touch in this area ^50,57,60,65^.

### When systematically exposed to delayed self-generated touches, responses in contralateral secondary somatosensory cortex become comparable for delayed and nondelayed self-generated touches, and their difference significantly decreases compared to the baseline

Next, we examined possible differences in the activation of the secondary somatosensory cortex between delayed and nondelayed self-generated touches in the *adaptation* session. We reasoned that if participants adapt to the sensorimotor delay when systematically exposed to delayed self-generated touches by recalibrating their internal model, then the nondelayed self-generated touches should elicit stronger somatosensory BOLD responses due to weaker sensorimotor predictions, and the delayed self-generated touches would elicit weaker somatosensory BOLD responses due to greater sensorimotor predictions. This should thus lead to elimination of the neural response difference in the secondary somatosensory cortex. Note that based our previous study ^46^ and the current behavioral data (see above), we did not expect a complete reversal of the sensory attenuation patterns for delayed and nondelayed trials after delay exposure.

Critically, there was no significant activation of the secondary somatosensory cortex (*p* < 0.001 uncorrected) or activation of any somatosensory area (*p* < 0.05 FWE-corrected) when we contrasted the delayed to the nondelayed self-generated touches in the *adaptation* session (*adaptation* < 0, **Supplementary Table S4**). Thus, the strongly significant activation of the right secondary somatosensory cortex observed in the *baseline* session (see above) was not observed in the *adaptation* session. To provide statistical support for the absence of the BOLD effect, we extracted the mean activity for delayed and nondelayed touches from the right secondary somatosensory cortex area we identified in the *baseline* session using an anatomical mask. For all runs in the *adaptation* session, as well as when considering each run separately, Bayesian statistics favored the null hypothesis (delayed ≤ nondelayed) over the alternative hypothesis (delayed > nondelayed) in the BOLD signal: all runs of *adaptation* session, *BF0-* = 9.266 (*adaptation* < 0); *early*, *BF0-* = 6.099 (*adaptation_early_* < 0); *middle*, *BF_0-_* = 9.553, (*adaptation_middle_* < 0); *late*, *BF_0-_* = 8.213 (*adaptation_late_* < 0) (**Supplementary Figure S4**). No region showed the opposite pattern of responses, either in the secondary somatosensory cortex (at *p* < 0.001 uncorrected) or anywhere in the brain (*p* < 0.05 FWE-corrected) (**Supplementary Table S5**). Moreover, we observed a significant cluster in the secondary somatosensory cortex when directly comparing the difference between delayed and nondelayed self-generated touches between the *adaptation* and the *baseline* sessions (*baseline* – (–*adaptation*), MNI: *x* = 58, *y* = -18, *z* = 22, *p* < 0.001 FWE-corrected) (**Figure 6, Supplementary Table S6**). This effect reflects significantly less secondary somatosensory cortex neural attenuation in the *adaptation* session compared to the *baseline* session for the nondelayed compared to the delayed touches.

**Figure 6.**
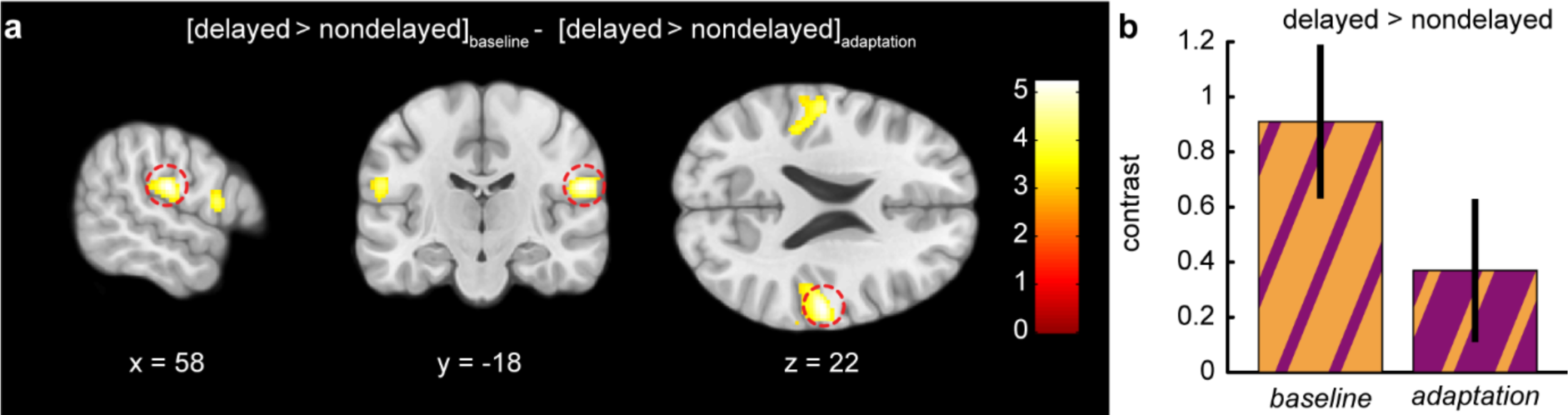
Systematic exposure to the sensorimotor delay led to significant changes in secondary somatosensory cortex activation for delayed versus nondelayed self-generated touches between the *baseline* and the *adaptation* sessions. (a) Sagittal, coronal, and axial views of the significant peak of activation (*p* < 0.05 FWE-corrected) located in the right secondary somatosensory cortex (parietal operculum; MNI: *x* = 58, *y* = -18, *z* = 22) during all trials of the *baseline* session compared to all trials of the *adaptation* session. The activation maps are rendered on the MNI-152 template brain and are displayed at an uncorrected threshold of *p* < 0.001. Red circles are centered over the main significant peak. The color bar indicates the values of the t-statistic. **(b)** Contrast estimates and 90% CI for the significant somatosensory peak in all runs of the *baseline* and *adaptation* session.

In conclusion, exposure to delays in the *adaptation* session led to a similar degree of neural activation for delayed and nondelayed self-touches in the secondary somatosensory cortex, *i.e.,* this area no longer distinguished between delayed and nondelayed self-generated touches. This provides neural evidence that the internal model has been recalibrated to a certain extent and that the delay adaptation modulates the neural somatosensory attenuation within the secondary somatosensory cortex.

### When systematically exposed to delayed self-generated touches, neural responses evoked by the nondelayed self-generated touches change over time in the ipsilateral cerebellum and gradually increase in the anterior cingulate areas

Our behavioral results and somatosensory cortical fMRI findings suggest that participants adapted to the delayed self-generated touches; yet, could we find evidence in support of the hypothesis that the cerebellum is involved in the recalibration of the internal model? In line with this, we observed a significant increase in the responses of the left anterior cerebellum (lobules I-IV) from the *early* to the *middle* runs that was subsequently reduced in the *late* run (*adaptation_middle_* > *adaptation_early_*, MNI: *x* = -16, *y* = -38, *z* = -26, *p* = 0.011 FWE-corrected) (**Figure 7a-b, Supplementary Table S7)**. No other significant cerebellar activations were observed from the *early* to the *late*, or from the *middle* to the *late*, or in all three contrasts in the opposite direction (*p* < 0.001 uncorrected).

**Figure 7.**
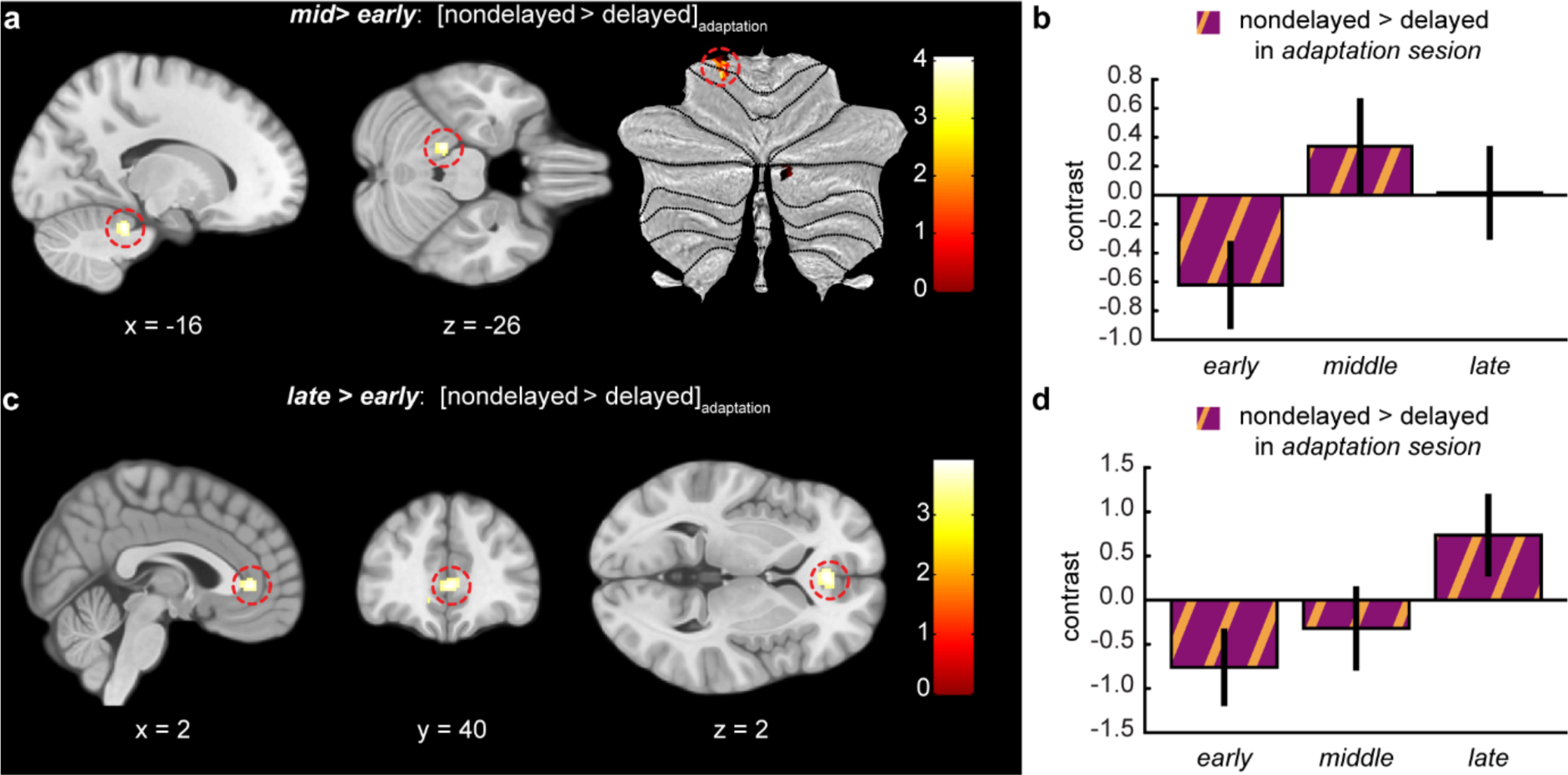
Cerebellar and cingulate activations during the nondelayed compared to the delayed self-generated touches across the exposure time in the *adaptation* session. (a) Sagittal and axial views and cerebellar flatmap of the significant peak of activation (*p* < 0.05 FWE-corrected) located in the left cerebellum (lobules IV/V; MNI: *x* = -16, *y* = -38, *z* = -26) that increased its activity from the *early* to the *middle* runs of the *adaptation* session. **(c)** Sagittal, coronal, and axial views of the significant peak of activation (*p* < 0.05 FWE-corrected) located in the anterior cingulate cortex (parietal operculum; MNI: *x* = -2, *y* = 40, *z* = 2) that increased its activity from the *early* to the *late* runs of the *adaptation* session. **(a, c)** The activation maps are rendered on the MNI-152 template brain and are displayed at an uncorrected threshold of *p* < 0.001. Red circles are centered over the main peak. The color bar indicates the values of the t-statistic. **(b, d)** Contrast estimates and 90% CI for the peaks displayed in **(a, c)** across all three runs of the *adaptation* session.

Finally, we theorized that after delay adaptation, the nondelayed self-generated touches should be considered as more unexpected by the brain. This might, in turn, trigger an internal cognitive conflict because unexpected nondelayed self-touch is inconsistent with a lifetime of previous knowledge of experiencing (nondelayed) self-touches as fully predictable. In support of this hypothesis, we observed a significant increase in the anterior cingulate cortex during the *late* compared to the *early* run (*adaptation_late_* > *adaptation_early_*, MNI: *x* = 2, *y* = 40, *z* = 2, *p* = 0.025 FWE-corrected; MNI: *x* = -2, *y* = 38, *z* = 2, *p* = 0.029 FWE-corrected) (**Figure 7c, Supplementary Table S8**), which was gradual along the exposure time (**Figure 7d**). There were no other significant ACC activations (*p* < 0.05 FWE-corrected) in the opposite direction: from the *early* to *middle*, from *middle* to *early*, from *late* to *middle* and from *late* to *early* (*p* < 0.001 uncorrected).

## Discussion

Prominent motor control theories ^6^ posit that self-generated touches are perceptually and neurally attenuated because they are predicted by the internal model ^46,53,57,59,61,67^. However, the fundamental question of how such central predictions are formed—based on the sensory experience of self-touch—and how the internal model is recalibrated in the brain has not been previously addressed. In this study, we utilized fMRI alongside a self-touch paradigm where participants underwent hundreds of trials with delayed tactile feedback (100 ms). This approach enabled us to pinpoint changes in the BOLD signal that reflect the neural recalibration of sensory attenuation and the internal model. Our study yielded three main novel findings. First, the neural attenuation response in the secondary somatosensory cortex, observed during natural (non-delayed) self-touch in this and numerous prior studies (Blakemore, Wolpert, et al. 1998; Shergill et al. 2013; Kilteni et al. 2020, 2023), vanished after exposure to delay. This suggests that somatosensory cortical attenuation is a dynamic process governed by the internal model’s central predictions. Second, the anterior cerebellar hemisphere demonstrated a time-varying neural activity pattern in line with the hypothesis of internal model recalibration within this subcortical region. Third, the anterior cingulate cortex exhibited changing activity patterns, suggesting the accumulation of a cognitive conflict signal as self-generated touches began to be expected as delayed. Taken together, these findings extend our understanding of the neuroplasticity that drives the dynamic recalibration of the internal model in self-touch and, in a broader sense, sensorimotor control.

In the behavioral runs, we replicated our previous results ^46^ showing that participants perceive their nondelayed self-generated touches as stronger when exposed to delayed (88%) compared to when exposed to nondelayed (88%) self-generated touches. This reduction in attenuation was present in the *late* phase of the exposure, which aligns with the timeline reported in our previous study ^46^. Earlier studies have shown that the systematic exposure to delays between a voluntary action and its visual, auditory, or somatosensory feedback induces a dynamic recalibration in the perceived timing: for example, the delayed sensory feedback is perceived as if occurring synchronously with the action, and/or the nondelayed sensory feedback as if occurring before the action ^33,38,105–110^. In line with this previous literature, the current results suggest that participants shifted the expected timing of their self-generated touches towards the delayed timing, reducing the attenuation of their natural nondelayed self-touches.

At the neural level, when participants were not exposed to the sensorimotor delay (*baseline* session), we observed stronger responses in the contralateral secondary somatosensory cortex during delayed compared to nondelayed self-generated touches. As said, this finding replicates previous studies that have described predictive neural attenuation of self-touch in this specific somatosensory area ^50,57,60,65^. In contrast, when participants were exposed to the sensorimotor delay (*adaptation* session), activity in the secondary somatosensory cortex activity was indistinguishable for delayed and nondelayed touches, and Bayesian statistics provided support for the absence of an effect. This observation is in line with our hypothesis that the neural responses to the nondelayed self-generated touches increase and the neural responses to delayed self-generated touches decrease over time, effectively minimizing their difference. Critically, a direct comparison with the *baseline* session revealed that a significant change in the activity of the secondary somatosensory cortex occurred due to the delay exposure; that is, the predictive attenuation of somatosensory processing in the secondary somatosensory cortex that was robustly present in the *baseline* session vanished in the *adaptation* session, which provides neural evidence for a recalibration of the internal model of self-touch. This pattern of findings is important as it suggests that the predictive attenuation of tactile input in the somatosensory cortex is a dynamic process that is subject to neuroplasticity and recalibration – in line with the remarkable capacity of the sensorimotor system to adapt to internal and external changes ^7,8,111^.

The behavioral responses showed signs of delay adaptation in the *late* run (**Figure 4**), but the neural somatosensory responses showed adaptation already from the *early* run. Although this may seem inconsistent, one should consider that the neural changes we observed in the *adaptation* session stem from changes affecting both delayed and nondelayed self-generated touches, while the behavioral responses assessed the perception of only the nondelayed self-generated touches. From a theoretical perspective, when participants are systematically exposed to a sensorimotor delay, they should update their internal model to predict the delayed touches and less so the nondelayed touches. Although both effects might be seen as two sides of the same coin – as a shift in expected temporal delay (Kilteni et a 2019) – and we have previously observed that the magnitudes of these effects were correlated (^46^, Experiment 1), they might not have the same time course. In our previous study (^46^, Experiment 2), we observed that changes in the perception of delayed self-generated touches occurred earlier than changes in the perception of nondelayed self-generated touches. That is, fewer exposure trials were needed to observe a significant decrease in the perceptual responses to delayed touches than the number of trials needed to observe a significant increase in the perceptual responses to nondelayed touches. Consequently, it might be that the observation of earlier changes in somatosensory activity in the *adaptation* session compared to the behavioral changes in sensory attenuation in the present study is due to the delayed touches becoming attenuated earlier, as reflected in the BOLD responses (i.e., delayed – nondelayed) as compared to the nondelayed touches that only show behavioral evidence of delay adaptation later.

However, if the activity change in the secondary somatosensory cortex reflects updates in somatosensory attenuation due to recalibration of the internal model, where is this recalibration implemented at the neural level? There is substantial agreement in the literature that the cerebellum is involved in implementing the internal model ^8,23,25,76,112–115^. The anatomy and morphology of the cerebellum make it particularly suited to implement the internal forward model ^12,25,71^ and learn through error signals ^116^. The major input to the cerebellum is conveyed through the cortico-ponto-cerebellar pathway, and cortical areas including motor, premotor and parietal areas form parallel loops with specific areas in the cerebellum (cerebrocerebellum) ^117,118^. In monkeys, the primary motor cortex (M1) sends strong projections to the cerebellar lobules V and VI ^117,119^, and studies focusing on pontocerebellar projections (*i.e.*, mossy fibers) provide animal experimental evidence that during voluntary limb movement, the mossy fiber activity is modulated prior to the movement onset but after M1 activity, putting forward the hypothesis that these projections convey the efference copy from M1 to the cerebellum ^71,73,120^. This together with the fact that the cerebellum receives proprioceptive and cutaneous feedback directly from the body and somatosensory input might be available also from the cortex (i.e., through the cerebrocerebellum) supported the view that the cerebrocerebellum might act as the internal forward model ^71,120^.

In line with this view and our hypothesis, we observed increased activity in the anterior cerebellum ipsilateral to the touched left hand in the *middle* run of the *adaptation* session compared to the *early* run, which then decreased in the *late* run. It is important to note that this cerebellar activity was localized in the hemispheres of lobules IV/V, which are part of the sensorimotor cerebellum that is connected to sensorimotor cortical areas and play a role in sensorimotor integration ^88,97,121^. Moreover, it is critical to mention that this increase was not caused by changes in how participants pressed the sensor with their right hand in the *middle* compared to the *early* or *late* runs, as our Bayesian control analysis provided decisive evidence in favor of comparable magnitude of press force between runs and sessions (*BF_01_* = 5×10^9^), and all forces were included as parametric modulators of the BOLD activity, meaning that we regressed out any possible force-related modulations of the fMRI signals. Thus, putative differences in muscular force between delayed and nondelayed trials cannot explain our findings ^86,87,122^. Thus, how does this increased activity relate to the internal model recalibration? Work in primates has shown that deep cerebellar neurons increase their activity in response to unexpected sensory feedback (Brooks et al., 2015; Brooks and Cullen, 2013) but this activity decreases as the new sensorimotor mapping is learned and returns to normal firing patterns (Brooks et al., 2015). Previous studies on motor learning have shown that cerebellar areas modulate their activity over the time course of learning new sensorimotor mappings as, for example, in visuomotor rotation ^123,124^, and this increased activity is prominent during the early phase of learning but gradually decrease as learning progresses ^125^. Moreover, earlier findings showed that inhibitory cerebellar transcranial magnetic stimulation prevents adaptation to delayed self-generated sounds ^37^. Considering the current finding and previous literature, we propose that the observed cerebellar activity reflects error-related activity that is higher in the *middle* run due to the accumulative exposure to the delay and that is subsequently resolved by the recalibration of the internal model for self-touch.

The anterior cingulate cortex showed gradually increasing activity to nondelayed self-generated touches during the *adaptation* session. The ACC has long been implicated in monitoring and resolving cognitive errors ^126^, detecting response conflict ^77–81^, computing prediction errors ^82^, and signaling the unexpectedness of the outcomes of an action ^83^. Activity in the ACC has been previously reported when investigating adaptation to temporal delays between voluntary taps and subsequently delivered light flashes ^33^. Specifically, Stetson and colleagues observed that – after adapting to delays between movement and sensory feedback – flashes presented without an injected delay and perceived as occurring *before* the participants’ movement was associated with activation of the anterior cingulate cortex and medial frontal cortex, interpreted by the authors as a cognitive conflict in perceiving that effects (flashes) precede the cause (movement). In line with this interpretation, we speculate that the gradual ACC activation to nondelayed self-generated touches in the present study reflects a cognitive conflict that builds up as unexpected nondelayed self-touch conflicts with a lifetime of previous experience of nondelayed self-touches as fully predictable. Note, that when unexpected delays are introduced in self-touch paradigms without preceding delay adaptation, activation of the ACC is typically not seen ^65^; indeed, we observed no such cingulate activity in our *baseline* session when sensory predictions were not met in the delayed self-touch trials. Therefore, we conclude that the anterior cingulate activity likely does not reflect a basic sensorimotor error signal that is used to recalibrate the internal model in the cerebellum but rather a higher-order cognitive conflict signal in the frontal association cortex related to expectation violation.

In the present study, we systematically exposed participants to the same constant delay between their movement and its somatosensory consequences (88% of the trials) to induce the delay adaptation and the recalibration of the internal model. We chose this learning “by repetition” as the most natural way of relearning the sensorimotor predictions in self-touch, similar to how we practice learning new sensorimotor skills and tools and the design of previous delay adaptation studies (*e.g.*, ^33,38,105,106, 108–110)^ and force-field and visuomotor adaptation studies (for reviews see ^8,23,127,128^). In relation to this method, it is important to stress that our results cannot be explained by differences in attention across sessions or by sensory habituation and neuronal adaptation to the repeated tactile stimuli. If this were the case, we should have observed the same patterns between the *adaptation* and the *baseline* session as these lasted the same duration, and the same patterns across all runs as all touches involved physically identical stimuli: forces of 2 N intensity, 250 ms duration, applied at the same region of skin on the pulp of the left index finger (**Supplementary Text S1, S2**). Neither the percentages of the delayed or nondelayed trials *per se* can account for our results, as in both sessions, we modeled the less frequent touches that had an identical number of appearances (*i.e.*, 12%) and differences between delayed and nondelayed self-generated touches have been previously observed in experimental contexts where predicted and unpredicted touches are equiprobable (50%-50%)^55,65^.

The present findings extend our knowledge regarding how the brain learns movement-related sensory predictions beyond earlier work in other species. Mice can learn to associate sounds with their movements, as if these sounds were the acoustic outcome of their actions ^129,130^. Once these self-generated sounds are learnt to occur at a certain time with systematic repetition, auditory cortical responses to sounds presented at that expected time are suppressed; in contrast, if the same sounds are delayed, suppression is smaller if not abolished ^18,130^. Additionally, moving larval zebrafish acutely react to perturbations in their optic flow (*i.e.*, visual reafference) in the opposite direction of their movement: for example, their bout lasts longer if their visual reafference lags. Interestingly, this initial acute reaction decreases substantially if the animals are systematically exposed to these sensorimotor lags, with the result that the duration of their bouts after persistent exposure become comparable to those of zebrafish that were never exposed to the perturbed visual reafference ^131^. Our results do not only extend these principal observations to the human brain but also reveal that the learning of movement-related predictions involves areas beyond the sensorimotor cortices (Schneider et al. 2018; Audette et al. 2022), including the sensorimotor cerebellum.

Here, we investigated the neural recalibration of self-touch, which unlike arbitrary mappings between movement and sensory effects (e.g., pressing a key and hearing a tone or seeing a Gabor patch), constitutes one of the most basic sensory-motor associations in human sensorimotor control and cognition, presumably established very early in development (or even in utero;^132^). The fact that the cerebellum and somatosensory cortex can flexibly re-learn the expected temporal delays in such a simple and extensively over-practiced sensorimotor task provides a particularly strong test case for the theory of internal models in sensorimotor control and predictive sensory attenuation. Second, self-touch is an important cue for bodily self-awareness and the distinction between bodily self and the external environment ^133–137^. Therefore, the findings that current somatosensory and cerebellar regions implement neural recalibration of self-touch can have broader implications for research into the plasticity of bodily self and how humans adapt to delays in brain-machine interfaces and haptic feedback in virtual reality applications.

In conclusion, the current neuroimaging results reveal a possible neural mechanism behind the temporal recalibration of sensorimotor predictions even in the very basic behavior of touching one’s own body. Together with earlier studies using animal models, the present study suggests that across species, sensory modalities, and tasks, the nervous system constantly recalibrates its internal models to keep sensorimotor predictions precise and in line with the current environmental statistics.

## Data availability

We do not have ethics approval to make the data publicly available.

## Author Contributions

**Konstantina Kilteni:** Conceptualization, Data curation, Formal Analysis, Funding acquisition, Investigation, Methodology, Software, Visualization, Writing – original draft, Writing – review & editing. **H. Henrik Ehrsson**: Conceptualization, Funding acquisition, Methodology, Supervision, Writing – review & editing.

## Acknowledgments

The authors would like to thank Christian Houborg, Sophie O Kane, Pius Kern and Sara Coppi for helping with the data collection.

