## Supplementary material for "Dynamic changes in somatosensory and cerebellar activity mediate temporal recalibration of self-touch"

**Text S1. Participants received comparable touches (test taps) and performed comparable presses (active taps) across all behavioral runs.**

We assessed whether participants received comparable touches (test taps) and performed comparable presses (active taps) across the 6 behavioral runs (**Figure S1)**. This was deemed necessary to ensure that any potential perceptual effects detected during the behavioral runs were not driven by the physical magnitude of the touch participants received on their left index finger or by the magnitude of the participants' presses of their right index finger.

To test this, we conducted a two-way RM ANOVA on the *test* taps with two within-subject factors: session (*adaptation* or *baseline*) and run (*early*, *middle*, *late*). Indeed, the ANOVA revealed a nonsignificant main effect of session ($F(1, 23) = 1.886$, $p = 0.183$), a nonsignificant main effect of run ($F(2, 46) = 2.101$, $p = 0.134$), and a nonsignificant session × run interaction ($F(2, 46) = 0.308$, $p = 0.737$) – this was strongly supported by a Bayesian two-way RM ANOVA favoring the absence of interaction ($BF_{01} = 14.839$). Similarly for the active taps, there was a nonsignificant main effect of session ($F(1, 23) = 0.315$, $p = 0.580$), a nonsignificant main effect of run ($F(1.544, 35.509) = 0.730$, $p = 0.\ 554$), and a nonsignificant session × run interaction ($F(2, 46) = 0.427$, $p = 0.655$) strongly supported by a Bayesian RM ANOVA ($BF_{01} = 37.995$).

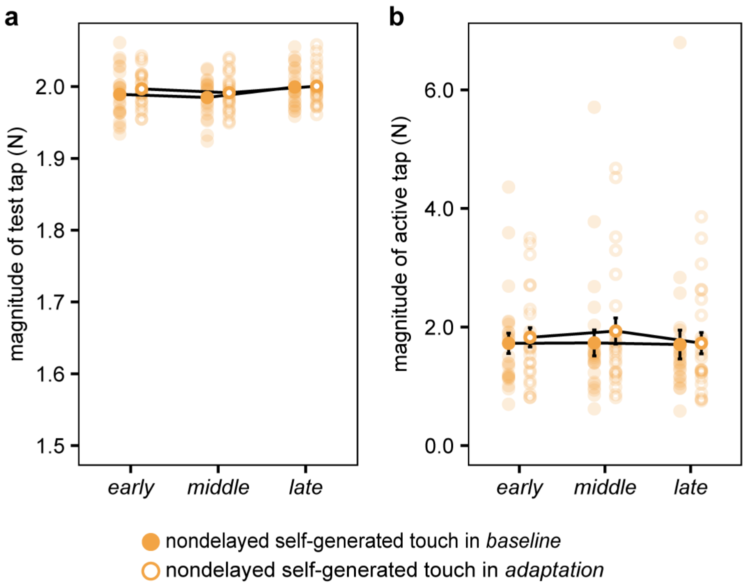


**Figure S1. Control analyses for the behavioral runs. (a-b)** Individual and group (mean ± s.e.m.) test (2N) and active taps per session and run. There were no statistically significant differences between any of the runs or sessions. Error bars represent the standard error of the mean. Note that the error bars of the magnitude of the test taps (2 N) are small since these forces were delivered by the electric motor.

**Text S2. Participants received comparable touches (test taps) and performed comparable presses (active taps) across all fMRI runs.**

Identical to the behavioral runs, we assessed whether participants received comparable touches (test taps) and performed comparable presses (active taps) across the 6 fMRI runs (**Figure S3)**. This was deemed necessary to ensure that any changes in BOLD activity during the fMRI runs were not driven by the physical magnitude of the touch participants received on their left index finger or by the magnitude of the participants' presses of their right index finger.

To test this, we conducted a three-way RM ANOVA on the *test* taps with three within-subject factors: session (*adaptation* or *baseline*), run (*early*, *middle*, *late*) and condition (nondelayed or delayed self-generated touches). The ANOVA revealed a nonsignificant main effect of session ($F(1, 23) = 2.532$, $p = 0.125$), a nonsignificant main effect of run ($F(2, 46) = 3.181$, $p = 0.051$), and a nonsignificant main effect of condition ($F(1, 23) = 0.003$, $p = 0.960$). All two-way interactions were nonsignificant: session × run ($F(2, 46) = 1.040$, $p = 0.361$), session × condition ($F(1, 23) = 0.622$, $p = 0.438$), run × condition ($F(2, 46) = 0.436$, $p = 0.649$). The three-way interaction session × run × condition was also nonsignificant ($F(2, 46) = 0.737$, $p = 0.484$) and a Bayesian RM ANOVA decisively supported the absence of the interaction effect ($BF_{01} = 1512.7$).

Similar for the *active* taps, a three-way RM ANOVA revealed a nonsignificant main effect of session ($F(1, 23) = 1.432$, $p = 0.341$), a nonsignificant main effect of run ($F(2, 46) = 0.058$, $p = 0.944$), and a nonsignificant main effect of condition ($F(1, 23) = 0.044$, $p = 0.835$). All two-way interactions were nonsignificant: session × run ($F(2, 46) = 0.722$, $p = 0.461$), session × condition ($F(1, 23) = 0.766$, $p = 0.391$), run × condition ($F(2, 46) = 0.758$, $p = 0.474$). Finally, the three-way interaction session × run × condition was nonsignificant ($F(2, 46) = 0.347$, $p = 0.709$) and a Bayesian RM ANOVA decisively supported the absence of the interaction effect ($BF_{01} = 5\times10^9$).

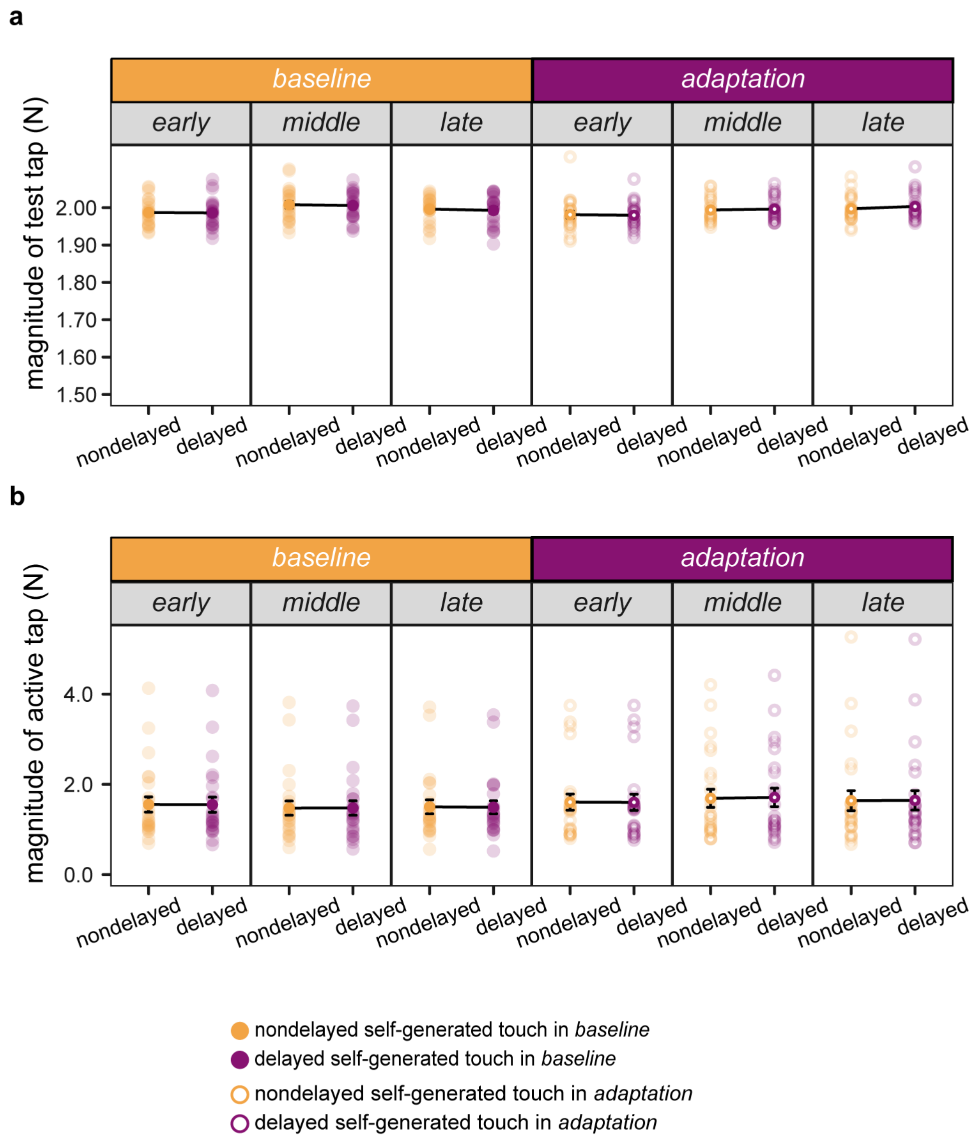


**Figure S2. Behavioral results from the fMRI runs.** (**a-b**) Individual and group (mean ± s.e.m.) test (2N) and active taps, per session and run. There were not any statistically significant differences between any of the runs or sessions. Error bars represent the standard error of the mean. Note that the error bars of the magnitude of the test taps (2 N) are small since these forces were delivered by the electric motor.

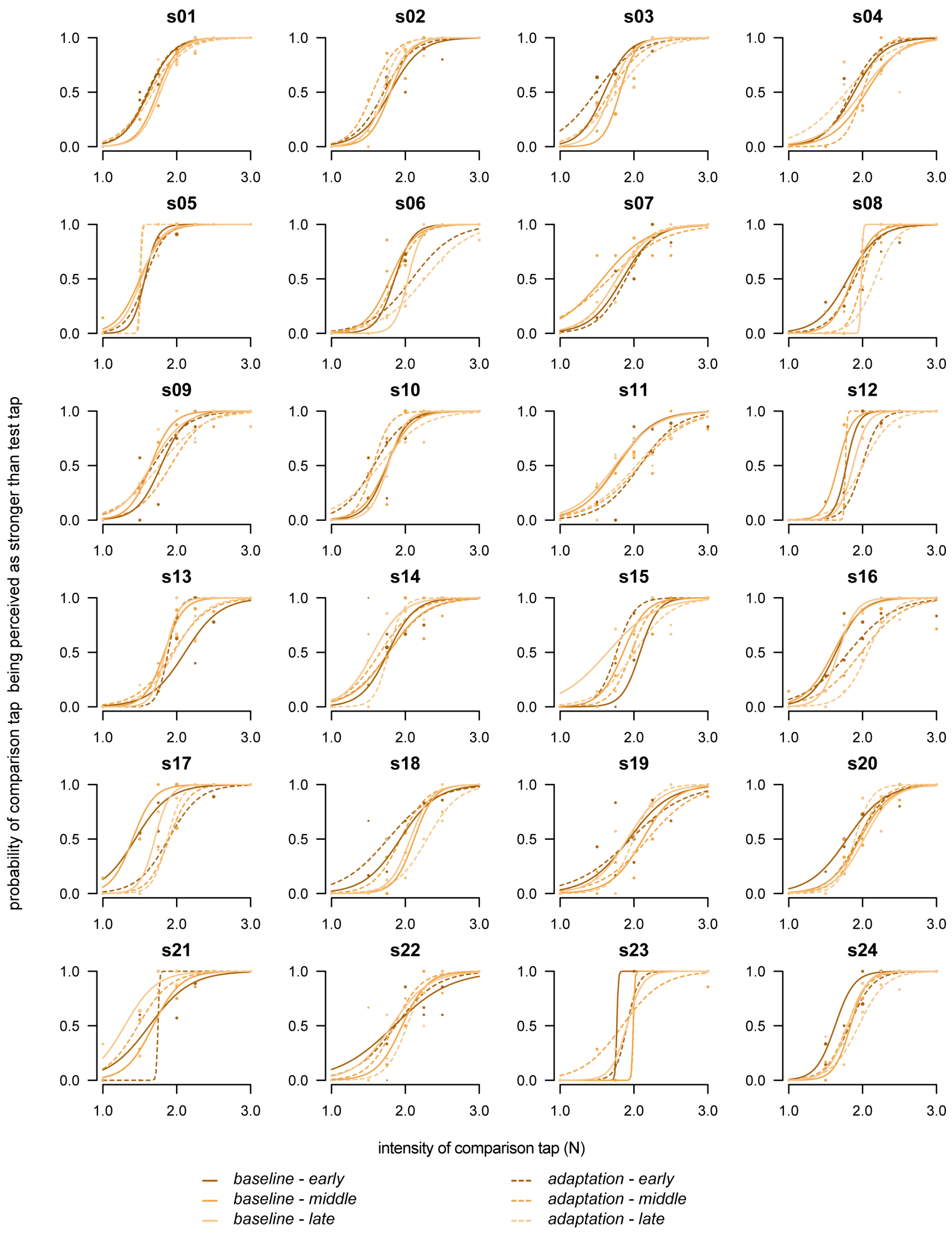


**Figure S3. Individual plots of the psychophysical task.** The marker size is proportional to the number of repetitions at that comparison tap intensity level. For all participants and conditions, the fitted model resulted in a McFadden's $R^2$ value ranging between 0.352 and 0.971.

**Table S1. Activations greater during the delayed compared to the nondelayed self-generated touches in the *baseline* session (all trials collapsed).** Only clusters with size greater than 4 voxels and peaks in grey matter are displayed.

| Brain region | Cluster size (voxels) | MNI coordinates (mm) | | | *z* | *p* |
|---|---|---|---|---|---|---|
| | | x | y | z | | |
| **R parietal operculum (SII)** | 252[1] | 54 | -20 | 22 | 5.39 | $p < 0.001$ FWE-corrected[*] |
| **R planum temporale (IPL)** | | 64 | -28 | 16 | 3.61 | $p < 0.001$ uncorrected |
| **R precentral gyrus** | 42 | 62 | 10 | 18 | 3.96 | $p < 0.001$ uncorrected |
| **L parietal operculum (SII)** | 207 | -44 | -30 | 20 | 3.84 | $p < 0.001$ uncorrected |
| **L parietal operculum (SII)** | | -50 | -24 | 22 | 3.78 | $p < 0.001$ uncorrected |
| **L supramarginal gyrus** | | -60 | -28 | 24 | 3.65 | $p < 0.001$ uncorrected |

[*] After small-volume correction. The peak was significant at the whole brain level too.
[1] The cluster size was 560 before corrections for multiple comparisons and was reduced to 252 after small-volume correction.

**Table S2. Activations during the delayed compared to the nondelayed self-generated touches that were greater in the *middle* than in the *early* run of the *baseline* session ($baseline_{middle}$ > $baseline_{early}$).** Only clusters with size greater than 4 voxels and peaks in grey matter are displayed.

| Brain region | Cluster size (voxels) | MNI coordinates (mm) | | | *z* | *p* |
|---|---|---|---|---|---|---|
| | | x | y | z | | |
| **R brainstem** | 45 | 6 | -46 | -54 | 3.72 | $p < 0.001$ uncorrected |
| **R superior frontal gyrus** | 27 | 12 | -2 | 70 | 3.57 | $p < 0.001$ uncorrected |

**Table S3. Activations during the delayed compared to the nondelayed self-generated touches that were greater in the *late* than in the *middle* run of the *baseline* session (*baseline*$_{late}$ > *baseline*$_{middle}$).** Only clusters with size greater than 4 voxels and peaks in grey matter are displayed.

| Brain region | Cluster size (voxels) | MNI coordinates (mm) | | | *z* | *p* |
|---|---|---|---|---|---|---|
| | | x | y | z | | |
| **R amygdala** | 49 | 28 | -12 | -12 | 3.62 | $p < 0.001$ uncorrected |
| **L supramarginal gyrus** | 4 | -64 | -42 | 24 | 3.37 | $p < 0.001$ uncorrected |

**Table S4. Activations greater during the delayed compared to the nondelayed self-generated touches in the *adaptation* session (all trials collapsed).** Only clusters with size greater than 4 voxels and peaks in grey matter are displayed.

| **Brain region** | **Cluster size (voxels)** | **MNI coordinates (mm)** | | | ***z*** | ***p*** |
|---|---|---|---|---|---|---|
| | | x | y | z | | |
| **R lateral occipital cortex** | 69 | 10 | -62 | 66 | 4.17 | $p < 0.001$ uncorrected |
| **R superior parietal lobule** | 129 | 30 | -42 | 68 | 3.79 | $p < 0.001$ uncorrected |
| | | 20 | -42 | 74 | 3.62 | $p < 0.001$ uncorrected |
| **R postcentral gyrus** | | 24 | -34 | 62 | 3.50 | $p < 0.001$ uncorrected |
| **R middle frontal gyrus** | 56 | 30 | 2 | 52 | 3.72 | $p < 0.001$ uncorrected |
| **L lateral occipital cortex** | 69 | -10 | -66 | 56 | 3.56 | $p < 0.001$ uncorrected |
| **L superior parietal lobule** | 27 | -22 | -44 | 62 | 3.53 | $p < 0.001$ uncorrected |
| **R hippocampus** | 39 | 20 | -24 | -14 | 3.49 | $p < 0.001$ uncorrected |
| **L lateral occipital cortex** | 11 | -18 | -62 | 42 | 3.32 | $p < 0.001$ uncorrected |
| **L postcentral gyrus** | 18 | -24 | -34 | 72 | 3.31 | $p < 0.001$ uncorrected |
| **L precuneous gyrus** | 6 | -6 | -56 | 66 | 3.15 | $p = 0.001$ uncorrected |

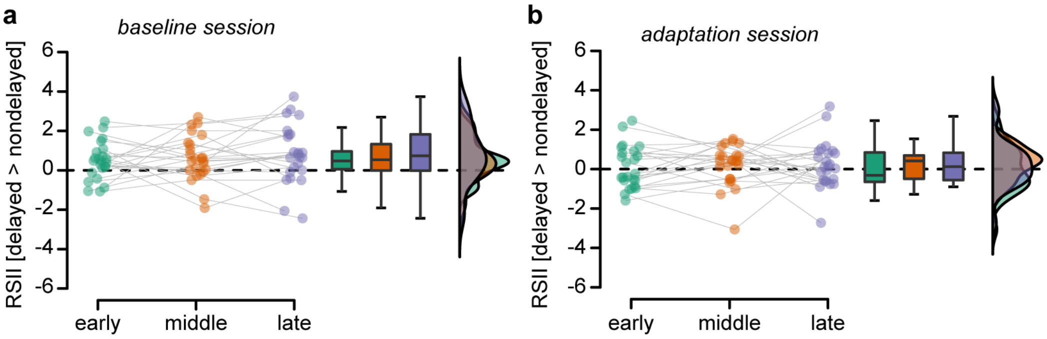


**Figure S4. Extracted activity from right SII (10 mm radius ROI) in each session and run.** Individual data, boxplots, and raincloud plots for the average activity within right SII extracted for all runs of the (a) *baseline* and (b) *adaptation* sessions. As seen, the differences were larger in the *baseline* than in the *adaptation* session. The black dashed line is centered at zero.

**Table S5. Activations greater during the nondelayed self-generated touches compared to the delayed self-generated touches in the *adaptation* session (all trials collapsed).** Only clusters with size greater than 4 voxels and peaks in grey matter are displayed.

| **Brain region** | **Cluster size (voxels)** | **MNI coordinates (mm)** | | | ***z*** | ***p*** |
|---|---|---|---|---|---|---|
| | | x | y | z | | |
| **L postcentral gyrus** | 5 | -56 | -16 | 22 | 3.29 | $p < 0.001$ uncorrected |
| **R superior temporal gyrus/supramarginal gyrus** | 4 | 66 | -32 | 16 | 3.24 | $p = 0.001$ uncorrected |

**Table S6. Activations greater during the delayed self-generated touches compared to the nondelayed self-generated touches in the *baseline* compared to the *adaptation* session (all trials collapsed).** Only clusters with size greater than 4 voxels and peaks in grey matter are displayed.

| **Brain region** | **Cluster size (voxels)** | **MNI coordinates (mm)** | | | ***z*** | ***p*** |
|---|---|---|---|---|---|---|
| | | x | y | z | | |
| **R parietal operculum (SII)** | 195[1] | 58 | -18 | 22 | 4.97 | $p < 0.001$ FWE-corrected[*] |
| **R superior temporal gyrus** | | 66 | -32 | 16 | 4.44 | $p < 0.001$ uncorrected |
| **R precentral gyrus** | 78 | 60 | 10 | 14 | 4.36 | $p < 0.001$ uncorrected |
| **L parietal operculum (SII)** | 208 | -56 | -18 | 22 | 4.09 | $p < 0.001$ uncorrected |
| **L parietal operculum (SII)** | | -44 | -32 | 22 | 3.66 | $p < 0.001$ uncorrected |
| **L supramarginal gyrus** | | -60 | -28 | 22 | 3.33 | $p < 0.001$ uncorrected |

[*] After small-volume correction. The peak was significant at the whole brain level too ($p$ = 0.006 FWE-corrected).

[1] The cluster size was 450 before corrections for multiple comparisons and was reduced to 195 after small-volume correction.

**Table S7. Activations greater during the nondelayed self-generated touches compared to the delayed self-generated touches in the *middle* compared to the *early* run of the *adaptation* session (*adaptation*$_{middle}$ > *adaptation*$_{early}$).** Only clusters with size greater than 4 voxels and peaks in grey matter are displayed.

| Brain region | Cluster size (voxels) | MNI coordinates (mm) | | | *z* | *p* |
|---|---|---|---|---|---|---|
| | | x | y | z | | |
| **L cerebellum IV/V** | 33[1] | -16 | -38 | -26 | 3.93 | $p = 0.011$ FWE-corrected[*] |
| **R cuneal cortex** | 9 | 22 | -72 | 20 | 3.52 | $p < 0.001$ uncorrected |
| **L hippocampus** | 15 | -28 | -38 | -6 | 3.38 | $p < 0.001$ uncorrected |
| **R cerebellum crus II** | 6 | 12 | -80 | -32 | 3.22 | $p = 0.001$ uncorrected |
| **L superior temporal gyrus** | 4 | -60 | -18 | 10 | 3.19 | $p = 0.001$ uncorrected |

[*] After small-volume correction.
[1] The cluster size was 52 before corrections for multiple comparisons and was reduced to 33 after small-volume correction.

**Table S8. Activations greater during the nondelayed self-generated touches compared to the delayed self-generated touches in the *late* compared to the *early* run of the *adaptation* session (*adaptation*$_{late}$>*adaptation*$_{early}$).** Only clusters with size greater than 4 voxels and peaks in grey matter are displayed.

| Brain region | Cluster size (voxels) | MNI coordinates (mm) | | | *z* | *p* |
|---|---|---|---|---|---|---|
| | | x | y | z | | |
| **R anterior cingulate gyrus** | 38[1] | 2 | 40 | 2 | 3.77 | $p = 0.025$ FWE-corrected[*] |
| **L anterior cingulate gyrus** | 23[1] | -2 | 38 | 2 | 3.72 | $p = 0.029$ FWE-corrected[*] |
| **R anterior cingulate gyrus** | 12 | 10 | 34 | -6 | 3.31 | $p < 0.001$ uncorrected |

[*] After small-volume correction.
[1] The cluster size was 78 before corrections for multiple comparisons with a peak at MNI: $x = 0$, $y = 38$, $z = 2$.
